# Reinstatement of emotional associations during human sleep: an intracranial EEG study

**DOI:** 10.1101/2022.06.24.497499

**Authors:** Guillaume Legendre, Laurence Bayer, Margitta Seeck, Laurent Spinelli, Sophie Schwartz, Virginie Sterpenich

## Abstract

The scientific literature suggests that emotional memories benefit from a privileged consolidation over neutral memories. This effect extends to consolidation processes that occur during sleep. Indeed, during sleep, a complex set of oscillations (namely slow-oscillations, theta rhythm and spindles) mediates the communication between brain regions involved in the long-term integration of memories. However, whether sleep oscillations may contribute to the reactivation and consolidation of emotional memories in humans is still unclear. Because non-invasive electroencephalography (EEG) has limited access to deep brain regions implicated in memory and emotion (e.g., hippocampus, amygdala, orbitofrontal cortex), here we recorded EEG signal from these brain regions using intracranial electrodes placed in medically-resistant epileptic patients in the context of presurgical investigation. During wakefulness, we presented the patients with emotional (i.e., humorous) vs emotionally neutral pictures paired with a sound. Then, we tested for the reinstatement of emotional-associations by delivering the sound during a subsequent period of sleep. We found that the reactivation of emotional (compared to neutral) memories during sleep enhanced slow-oscillation and spindle activity in the orbitofrontal cortex, paralleled with an increase in theta connectivity between the hippocampus and the orbitofrontal cortex.

In addition, we observed that the theta response to emotional memories reactivated at subsequent wake was different than for neutral memories, suggesting a change in memory traces with targeted memory reactivation. These data suggest that consolidation of emotional events during sleep is due to a larger expression of sleep features (in the slow-oscillation, theta and sigma frequency bands) and that the mechanisms of brain plasticity also take place in emotional brain regions during NREM sleep.

## Introduction

The most mundane, even unnoticed, sensory stimulation can elicit strong emotions. In the novel “In search of Lost Time” (Proust, 1922), the main character, Swann, is suddenly submerged by an intense emotion when eating a madeleine dipped in lime blossom tea: the taste of the madeleine had brought back joyful childhood memories to life. The reinstatement of emotions (positive or negative) linked to a specific past situation or event is an adaptive mechanism, which guides appropriate behavioral responses by signaling shared properties with the current situation or event (Nesse & Ellsworth, 2009). Moreover, stimuli or events with an associated emotional value are prioritized by memory consolidation mechanisms (LaBar & Cabeza, 2006). Such emotion-related memory benefit has been shown to be particularly relevant for consolidation processes occurring during sleep (Girardeau et al., 2009; Lipinska et al., 2019; Singer & Frank, 2009; Sterpenich et al., 2014; Trouche et al., 2020). For instance, after the encoding of novel information, a single night of sleep was found to favor the strengthening of emotional compared to neutral memory traces (P. Hu et al., 2006; Sterpenich et al., 2007, 2014; Wagner et al., 2006), by privileging the spontaneous neural replay of emotional information (Sterpenich et al., 2021). Igloi et al. (2015) reported that even a short nap can promote the long-term consolidation of emotional memory associations. In this same study, memory performance for emotional memories correlated with increased NREM sleep spindles, whose role in neural plasticity is well established (Klinzing et al., 2019). Thus, sleep may offer a permissive condition for the coordinated replay of memory traces in the hippocampus and thalamo-cortical neural networks (Rasch & Born, 2013). Such replay likely takes place during hippocampal sharpwave ripples, i.e. fast oscillations (80-200 Hz; Axmacher et al., 2008; de Lavilléon et al., 2015; Ego-Stengel & Wilson, 2010; Girardeau et al., 2009). Sharp-wave ripples generation seems potentiated during up-phases of the sleep spindles (12-15 Hz), while spindles themselves occur predominantly during the up-phase of slow-oscillations (<1 Hz) during NREM sleep (Clemens et al., 2007; Klinzing et al., 2016; Staresina et al., 2015; Takeuchi et al., 2016). Moreover, in rodents, functional interactions between the hippocampus and the ventral striatum are enhanced during sharp-wave ripples and considered to strengthen the association between memories and their affective (here emotional) value (Lansink et al., 2009). Together, these data suggest that oscillatory activity during sleep orchestrates neural reactivations across distributed brain networks. Whether and how the emotional aspect of a memory trace is reinstated in memory and emotion-related regions during human sleep is less well understood (see Sterpenich 2021).

In animals and humans, memory consolidation during sleep may be experimentally manipulated by presenting a sensory cue that had previously been associated with a particular memory element; a procedure also called “targeted memory reactivation” (TMR; Oudiette & Paller, 2013). Using TMR, the reactivation in sleep can be biased toward replaying a preselected memory element or context (Antony et al., 2012; X. Hu et al., 2015, 2020; Oudiette & Paller, 2013; Rasch et al., 2007) and influence the response of cortical circuits in which the memory is encoded on the next day (Bendor & Wilson, 2012), in particular when the memory is emotional (Sterpenich et al., 2014). In humans, performance at retrieval was found to correlate with TMR-induced sleep oscillations (such as spindles and slow-waves; Bar et al., 2020; Cairney et al., 2014, 2018; Creery et al., 2015; Lehmann et al., 2016), thus suggesting that TMR operates by hijacking endogenous plasticity mechanisms pertaining to sleep (see above). How does TMR influence temporally-organized neural oscillations across distributed brain regions is still unresolved. During wakefulness, the successful retrieval of memories was found to implicate theta neural synchronization between the hippocampus, the striatum and the neocortex (Herweg et al., 2016; Kaplan et al., 2014). It is thus plausible, yet not established, that TMR relies on a similar mechanism to foster the reactivation of emotional memory traces during NREM sleep.

Based on the observations reported above, we tested two main hypotheses pertaining to emotion memory reprocessing in human sleep. While sleep favors memory replay across hippocampal and thalamo-cortical networks, we propose that (i) the emotional dimension of the original event is reinstated across emotional brain networks, and that (ii) the reactivation of emotional memories during sleep implicates a functional interaction between emotional and memory systems, in particular via theta oscillations.

Here, we directly addressed these questions by analyzing intracranial EEG data recorded in 11 epileptic patients, selected based on their electrodes’ implantation sites. Specifically, we focused on three relevant brain regions of interest regarding our main hypotheses: the hippocampus for its documented contribution to sleep-related memory replay (Rasch & Born, 2013), as well as the amygdala and the orbitofrontal cortex (OFC) for their implication in emotion and more specifically humor processing (Bekinschtein et al., 2011; Chan, Hsu, & Chou, 2018; Chan, Hsu, Liao, et al., 2018). We used a TMR paradigm in which one auditory cue was paired with emotional (humorous) pictures while another auditory cue was paired with neutral (non-emotional) pictures. We then investigated the neural responses to these sounds when they were subsequently played during sleep. We used humorous stimuli because they elicit strong positive emotions, and recruit the mesolimbic reward system (Bekinschtein et al., 2011; Chan, Hsu, & Chou, 2018; Mobbs et al., 2003; Watson et al., 2007), which is involved in the spontaneous replay of rewarded events together with the hippocampus during sleep in rodents and humans (Lansink et al., 2009; Sterpenich et al., 2021). Moreover, positive pictures are well suited for testing patients in acute neurological conditions (here epileptic patients with implanted electrodes; see also Schwartz et al., 2008).

We found that the sound initially paired with emotional pictures (compared to the sound associated with neutral pictures) bolstered the generation of sleep oscillations typically involved in memory consolidation in emotional networks and induce a dialogue between emotion- and memory-related brain regions.

## Methods

### Patients

Fifteen patients with medication-resistant epilepsy were included in this study. They were tested while they had depth EEG electrodes implanted, as part of a standard presurgical procedure. Patients had normal or corrected-to-normal vision, and normal audition. The experimental procedure was approved by the Cantonal Ethics Commission for Research on Human Beings and all patients provided written informed consent. Four patients were rejected of the pool due to inaccurate electrode placement, corrupted signal or inaccurate markers timings. 6 out of the 11 patients achieved sustained sleep (reaching N3 sleep stage) during a nap in the medical environment, while 5 patients remained fully awake (see below). Therefore, we separated the dataset into two groups: 6 patients were included in a Sleep group and 5 patients were included in a Wake group. Note that the analysis implicating the OFC was performed on 5 out of the 6 patients from the Sleep group (one patient had no implantation in the frontal cortex). We would like to stress that the small patient sample is due to restriction related to electrode implantation sites and to our very strict handling of the data, especially regarding sleep and EEG data quality. Critically, for all the reported analyses, we applied the most stringent statistical methods recommended for small samples.

### Stimuli

#### Pictures

For the experiment, 90 humorous and 90 neutral pictures were selected from several database of stimuli and from internet matched for visual complexity and content (objects, characters, animals and actions depicted), as well as for mean luminance. Valence and intensity of emotion of pictures were scored by 12 external participants to validate the categorization in humorous and neutral pictures. For each patient and in each picture category (humorous, neutral), 60 pictures were randomly selected to appear both in the conditioning and memory test phase, while the remaining 30 pictures appeared only during the memory test phase as new images (see below). Note that the first participant saw 90 pictures per category in the conditioning phase and 120 in the memory phase) and, because this initial overloading of information was determined detrimental to the patient’s performance, the number of pictures was reduced for all subsequently tested patients.

#### Auditory stimuli

We created three different 600 ms pure tone sounds, respectively paired with humorous pictures, neutral pictures, and no picture (see Experimental paradigm below), and whose frequency was modulated in time as follows. One pure tone with a quadratically increasing frequency from 330 to 950 Hz, one pure tone with a quadratically decreasing frequency from 770 to 150 Hz, and one pure tone increasing quadratically from 220 to 880 Hz during 300 ms then decreasing back quadratically to 220 Hz during the last 300 ms. Sounds were tapered by a Gaussian window of the same length than the sounds (600 ms; “gausswin” matlab function) to avoid sharp intensity transition that could lead to awakening. Note that the sound not paired with pictures was not investigated in the present study as many publications comparing paired and non-paired sounds in TMR protocols already exists (see (Schouten et al., 2017) for a review). Sounds were delivered via speakers installed in patient’s room and controlled by the stimulation computer. Sound intensity was set with the patient so that a few sample sounds were clearly hearable during the tasks at wake. Before the nap, the sound intensity was decreased to be listenable but to avoid awakening and adjusted for each patient (see below).

### Experimental paradigm

The experiment was divided into four phases occurring sequentially just after lunch (between 12:00 and 18:00; Fig. 1A): 1) Sound exposure, 2) Conditioning, 3) Reactivation, and 4) Memory test.

**Figure 1.**
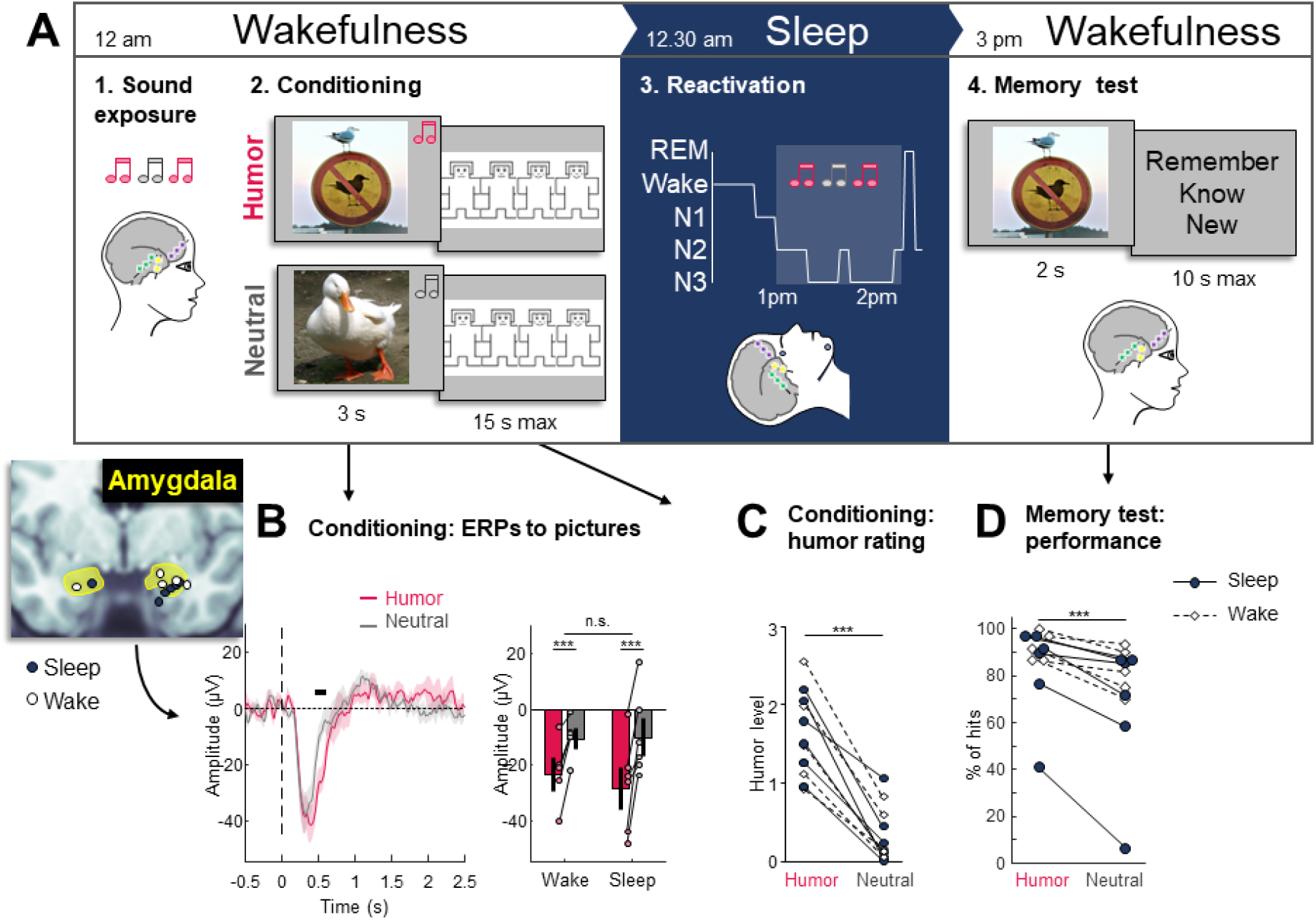
Main protocol, emotional response during encoding and behavior performances. (A) The graph summarizes the typical course of the experiment for a patient. At wakefulness, the patient is first exposed to sounds. Then, the conditioning procedure pairs one sound with humoristic pictures and another sound with neutral pictures. After that, the patient is offered a nap. The auditory cues are played randomly as soon as signs of NREM sleep appear. After the nap, the patient performs a memory test and has to dissociate between old pictures mixed with new ones. (B) The left subpanel depicts the position of the amygdala electrode of each patient of the sleep group (dark blue) and wake group (white). The middle subpanel shows the recorded ERP to neutral (gray) and humoristic (fuchsia) pictures in the amygdala for all the patients (N=11). The timepoints of significant difference between ERP to emotional and neutral pictures were averaged and reported in the right subpanel. (C) Mean humor rating of neutral and humorous pictures for each patient (small circles) in the sleep (dark blue) and wake (white) groups. (D) Performance of each patient on the memory task for humoristic and neutral pictures (small circles). The performance was computed as the number of hits (i.e. the number of familiar and remembered pictures recognized from the conditioning phase of the experiment). The average across participants within each category of pictures is depicted by a small cross on the side.

The goal of the **Sound exposure** phase was to check that the responses of the key brain regions (OFC, amygdala and hippocampus) to each of the three distinct sounds did not differ significantly. A sequence of sixty sounds (20 repetition of each sound type, randomly shuffled) was played with a jitter between 1800 to 2000 ms (A random integer between 1800 and 2000 was drawn from a uniform distribution) between sounds.

Then, patients underwent the **Conditioning phase** during which one neutral sound was paired with humorous pictures while another neutral sound was paired with neutral pictures. Specifically, 60 humorous and 60 neutral pictures were displayed centrally on a screen, organized in blocks of 6 pictures of the same type (either six humorous or six neutral pictures, pseudorandom order of blocks to avoid succession of 3 blocks of the same condition), and paired with their corresponding sound (see above, Stimuli section). For each trial, a fixation cross was displayed during 1 s, immediately followed by the presentation of a picture, shown during 2000 ms. Between 500 to 800 ms (A random integer between 500 and 800 was drawn from a uniform distribution each time to compute the length of the jitter in ms) after the onset of the picture, the sound corresponding to the picture type (i.e. humorous or neutral) was played during 600 ms. After the picture disappeared, a screen with 4 Manikins symbolizing 4 different levels of humor (from 0 –neutral, to 3 – very funny) were displayed and the patient rated how funny he/she found the picture by selecting one of the Manikins (15 s max, Fig. 1A). A pause of 4 to 15 s (A random number was drawn from a gaussian distribution with a mean of 0 and a standard deviation of 3, its absolute value was computed, it was rounded at the inferior and was added to 4 to compute the length of the pause in s) was inserted after every block, and a self-paced longer pause was allowed after blocks 7 and 14.

After the Conditioning phase, the preparation for the **Reactivation phase** started by placing surface electrodes for sleep monitoring: two electromyography (EMG) electrodes on the chin, two electrooculography (EOG) electrodes above and on the left of the left eye and below and on the right of the right eye, and two EEG electrodes on Fp1 and Fp2 positions when the head bandage allowed it. Then, the volume of the auditory cues delivered by a computer was adjusted to be barely hearable by the patient (to avoid waking him/her up during the nap session). The patient was then invited to rest and fall asleep, and the experimenters left the room. During the Reactivation phase, the experimenters monitored the online recordings and, as soon as stable NREM2 (K-complex or spindles) was detected, they started the presentation of the auditory cues. Sounds were triggered manually from the control room with a minimum delay of 3 s between two consecutive sounds. Every time the patient showed signs of awakenings, sound delivery was stopped, and resumed as soon as stable NREM2 was reached again. On average, the delay between sounds was of 4.19 seconds across participants, and the three types of sounds (paired with humorous pictures, paired with neutral pictures, and not paired) were presented randomly. The maximum number of stimulations was limited to 100 per sound type and was 54.58±29.45 per sound type on average (mean±SD; Number of “humor” cues = 54.36±33.63, Number of “neutral” cues = 54.09±26.42, Number of “new” cues = 55.27±30.10).

Following the nap, once the patient was fully awake and ready to continue, we proceeded to the **Test phase**, during which pictures and sounds were presented one at a time and in a fully random order. Every 10 trials, there was a short pause of 4 to 15 s (A random number was drawn from a gaussian distribution with a mean of 0 and a standard deviation of 3, its absolute value was computed, it was rounded at the inferior and was added to 4 to compute the length of the pause in s), and 3 longer self-paced pauses were allowed after trials 40, 80 and 120 (i.e. presentation of a sound or a picture). During a picture trial, a fixation cross was first displayed during 1 s, then the picture appeared for 2 s, followed by a third screen prompting a memory judgment (Fig. 1A). The patient had 10 s to answer by pressing a key whether he/she remembered the picture with vivid details from the encoding part (“remember”), whether he/she had the feeling of having seen the picture in the encoding part but couldn’t remember it in details (“know”), or whether it was a new picture, not seen in the encoding (“new”). Over a total of 180 pictures, 90 humorous and 90 neutral pictures were presented; each set was composed of 60 pictures seen at encoding and 30 new pictures (except for the patient 1 that showed 240 pictures, including 180 old (90 hum and 90 neu) and 90 new (45 hum and 45 neu) pictures). During a sound trial, a fixation cross appeared for 2.6 s, and the sound was played after 1 s. No response was required from the patient on the sound trials. There were 90 sound trials composed of the three sounds repeated 30 times each. Every 10 trials, there was a short pause of 4 to 15 s (A random number was drawn from a gaussian distribution with a mean of 0 and a standard deviation of 3, its absolute value was computed, it was rounded at the inferior and was added to 4 to compute the length of the pause in s), and a longer self-paced pause was allowed after every 40 trials (i.e. presentation of a sound or a picture).

### Electrophysiological recordings

#### Sleep data

Sleep during the nap session was scored by three experienced scorers according to the AASM rules (Iber et al., 2007) using EMG, EOG, and EEG signal from scalp electrodes (when available) or from depth electrodes close to the scalp.

#### Intracranial data acquisition

Stereo-EEG (SEEG) were recorded continuously during all phases of the experiment using depth electrode and linear shafts (Ad-Tech Medical, Racine, WI, or Dixi Medical, Chaudefontaine, France) and were then digitalized (CeeGraph, Natus Bio-logic Systems Corp. and then SystemPlus, Micromed). Electrodes were all referenced to a common scalp electrode localized on Cz position for bilateral implantation or contralateral C3 or C4 position for unilateral implantation. EEG data were recorded at a sampling rate of 1024 or 2048 Hz, and downsampled offline at a common sampling rate of 512 Hz. Offline signal processing was conducted with Fieldtrip functions (for downsampling and filtering; Oostenveld et al., 2010) and custom Matlab scripts.

#### Intracranial electrode selection

For each patient, a pre-implantation anatomical structural T1 (3T Siemens scanner; 1mm voxels) and a post-implantation CT scan were acquired. Pre-implantation MRI scans were first segmented using FreeSurfer (Reuter et al., 2012) and post-implantation CT scans were then co-registered using routines of FSL software (Jenkinson et al., 2012). Electrode localization and position normalization were performed according to the guidelines of the pipeline iELVis (Groppe et al., 2017). Specifically, electrodes contacts were localized using BioImage Suite (Papademetris et al., 2006) by thresholding the intensity of CT scans and finding the center of gravity of artifacts of high intensity caused by electrodes. T1 MRI scans were then normalized based on FsAverage brain template of FreeSurfer. Depth and surface electrode positions were normalized following two different techniques. Depth electrodes were normalized using the standard deformation field. To correct for brain shifts due to insertion of electrodes between pia and grey matter, surface electrode positions were normalized by inflating the native cortical surface up to a sphere and shrinking it back to the normalized brain cortical surface as detailed in Groppe et al. (2017). For each patient, three electrodes were selected, when possible, one in or close to the amygdala, one in or close to the hippocampus, and one in or close to the OFC (see Introduction section). If several electrodes were within the region of interest, we chose the electrode responding with the highest ERP amplitude to pictures (humorous and neutral together).

### EEG preprocessing

The EEG data were analyzed in terms of the amplitude (event-related potentials, ERPs), time-frequency power modulations and connectivity in response to the stimuli presented. We first applied three notch filters at 50, 100, and 150 Hz to the continuous EEG to remove line noise and bandpass-filtered the signal between 0.1 and 45 Hz using a 1st order two-pass butterworth filter. From the filtered continuous EEG recordings in each experimental phase, and for each type of stimulation (sound or picture), we extracted 5 s long epochs around each stimulus onset (from -2 to 3 s). The epoched signal was then downsampled once again at 128 Hz to reduce computation time. For each participant, we computed the standard deviation of all epochs recorded from the most artefacted brain region (i.e., the hippocampus). Within each phase, we discarded the epochs with a standard deviation exceeding 1.5 times the interquartile range of the distribution of standard deviation of epochs resulting in the rejection of 2.57±1.81% of all epochs (mean±SD). To extract time-frequency spectra, we convoluted the filtered continuous EEG signal with Morlet wavelets, with central frequencies 0.75 Hz and then from 1 to 30 Hz with steps of 1 Hz. The timespan of wavelets was computed so that the Full Width at Half Maximum (FWHM) was 1.5 s for 0.75 Hz wavelet then decreasing logarithmically from 1 s to 150 ms for 1 to 30 Hz wavelet respectively (Cohen, 2019). Then, the signal was epoched from -2 to 5 s to stimulus onset, artefacted epochs found with previous automatic rejection were discarded. Large epochs (−2 to 5 s) were extracted to subsequently compute the time course of connectivity with a sliding time window (see below) but only data from -1 to 2.5 s were considered for analysis, statistics and display of induced power and connectivity in the present study.

#### Power analysis

We computed the instantaneous power as 2 times the modulus of the complex number power 2. To correct for power difference between frequencies (1/f power law) and exponential growth of power, the instantaneous power was normalized with decibel correction by applying a tenth log-transform to the instantaneous power and multiply it by 10. Then, baseline correction was applied for each epoch by averaging the signal from -1 sec to stimulus onset and subtract this baseline value from the remaining time series.

The longer baseline (compared to ERPs) was set to account for slow fluctuations of power for low frequencies. To analyze SO activity, the power at the 0.75Hz was investigated (Achermann & Borbély, 1997). We then averaged the baseline-corrected time series across frequencies from 4 to 8 Hz (theta rhythm) and 12 to 15 Hz (sigma rhythm).

#### Connectivity analyses

We computed instantaneous phase as the argument function of the complex number. To test whether the reactivation of emotionally-positive associations implicated a change in the functional dialog between hippocampus and OFC (see Introduction section), we assessed the phase synchrony between both brain regions using the weighted Phase Locking Index (wPLI) approach as described in Vinck et al., (2011; see also Lachaux et al., 1999), and which is presumably insensitive to volume conduction. Specifically, a wPLI was computed for each time point at each frequency using a sliding window of 1.5 s. For each participant, the wPLI was baseline-corrected by subtracting the mean wPLI from -0.5 s to stimulus onset, and time series of wPLIs were averaged in the theta frequency range (4 to 8 Hz, see Introduction section).

### Statistics

Mean ± standard deviation of the sample is presented in the text unless stated otherwise. To account for small samples with unknown distributions, statistical testing was performed using a bootstrap resampling method. Both significance and confidence intervals were estimated using percentile method for one-sample testing, and studentized boostrap-t for two-samples testing. All statistical tests were performed two-sided and with 50000 resamplings (B=50000) or the exact number of possible resampling if it was below 50000 resamplings. Statistical tests were done with homemade scripts based on methods described by Efron & Tibshirani (1994). To analyze time-series, the boostrap procedure was applied at each time point, the family-wise error rate was controlled with Bonferroni correction by multiplying each p-value by the number of time points between 0 s and 2.5 s post-stimulus onset (N=321), and only statistical difference that lied on at least 10 consecutive time points were considered.

## Results

Behaviorally, during encoding, all patients rated the preselected humorous pictures (see Methods) as funnier than neutral pictures when considering both Sleep And Wake participants together (Fig. 1C; see also Table S1, first row; difference humoristic vs neutral on a scale from 0 – neutral to 3 – very funny: 1.27 ±0.405; bootstrap resampling with N=11, CI=[1.05,1.50], p<0.001, d=3.138) and no significant difference between the two groups was observed (difference between Wake and Sleep groups on the contrast Humoristic vs Neutral pictures: -0.0817±0.425; 2-samples bootstrap resampling with N=11; z=0.051; p=0.694; CI=[0.402,-0.741]; d=-0.192). During the memory test after the resting period, humorous pictures were better remembered than the neutral pictures (Fig. 1D; see also Table S1, 2^nd^ row; Sleep and Wake Groups together; Difference in percent of hits: 13.5±9.07; bootstrap resampling with N=11; z=5.041; p<0.001; ci=[8.81,19.3], d=1.489) and no significant difference was observed between groups (Sleep vs Wake Groups on contrast humoristic vs neutral pictures; percent of hits: -5.83±9.01; Bootstrap resampling with N=11; z=0.065; p=0.239; CI=[3.44,-19.1]; d=-0.647).

### Neural responses to sounds and pictures prior to sleep

We checked whether neural responses to the three distinct sounds differed at wake. During the sound exposure phase, ERPs from the amygdala, hippocampus, and OFC did not disclose significant differential responses to the two sounds (Fig. S1), suggesting that they are perceived similarly before the emotional conditioning. During the conditioning phase, both emotional and neutral pictures evoked robust and large negative wave in the amygdala, hippocampus and OFC (Fig. S2). Moreover, emotional pictures (compared to neutral pictures) appeared to trigger a sustained and significant negativity from 438 ms to 617 ms after picture onset in the amygdala, when considering both Sleep and Wake participants together (Fig. 1B; mean difference ±SD: -15.560 ±8.416 µV; bootstrap resampling with N=11; z=-6.408; p<0.001; CI=[-20.486,-11.028]; d=-1.85). Within the cluster of significant time points, responses to funny and neutral pictures differed significantly for both the Sleep group (−18.198 ±10.093 µV; bootstrap resampling with N=6; z=-4.863; p<0.001; CI=[-25.374,-11.006]; d=-1.80) and the Wake group (−12.395 ±5.175 µV; bootstrap resampling with N=5; z=-5.987; p<0.001; CI=[-16.111,-8.081]; d=-2.40), with no between group difference (5.803 ±8.276 µV; N=11; z=-0.005; p=0.249; CI=[16.019,-4.625]; d=0.70).

### Neural responses to auditory cues during sleep

After the conditioning phase, we recorded the activity of the amygdala, hippocampus and OFC when auditory cues were played during sleep (i.e. in the Sleep group). Using power spectrum analysis, we found enhanced SO, theta and sigma activity in OFC (Fig. 2A) in response to the auditory cue associated with humorous pictures compared to the sound paired with neutral pictures (Fig. 2B and 2C). We next analyzed the time course of these frequency bands. We observed that these frequency modulations disclosed a specific temporal ordering of activation, with SO increase arising first (largest cluster where difference is significant from 289 to 453 ms; mean difference ±SD: 1.475 ±0.946 dB; bootstrap resampling with N=5; z=3.899; p<0.001; CI=[0.721,2.182]; d=1.56), followed by theta power increase (largest cluster from 469 to 953 ms; 1.299 ±1.045 dB; bootstrap resampling with N=5; z=3.107; p<0.001; CI=[0.626,2.237]; d=1.24), and then sigma power increase (largest cluster from 1220 to 1570 ms; 1.820 ±1.255 dB; bootstrap resampling with N=5; z=3.626; p<0.001; CI=[0.888,2.878]; d=1.45). No significant difference between conditions for these 3 frequency bands were founded in the amygdala and in the hippocampus (Fig S3).

**Figure 2.**
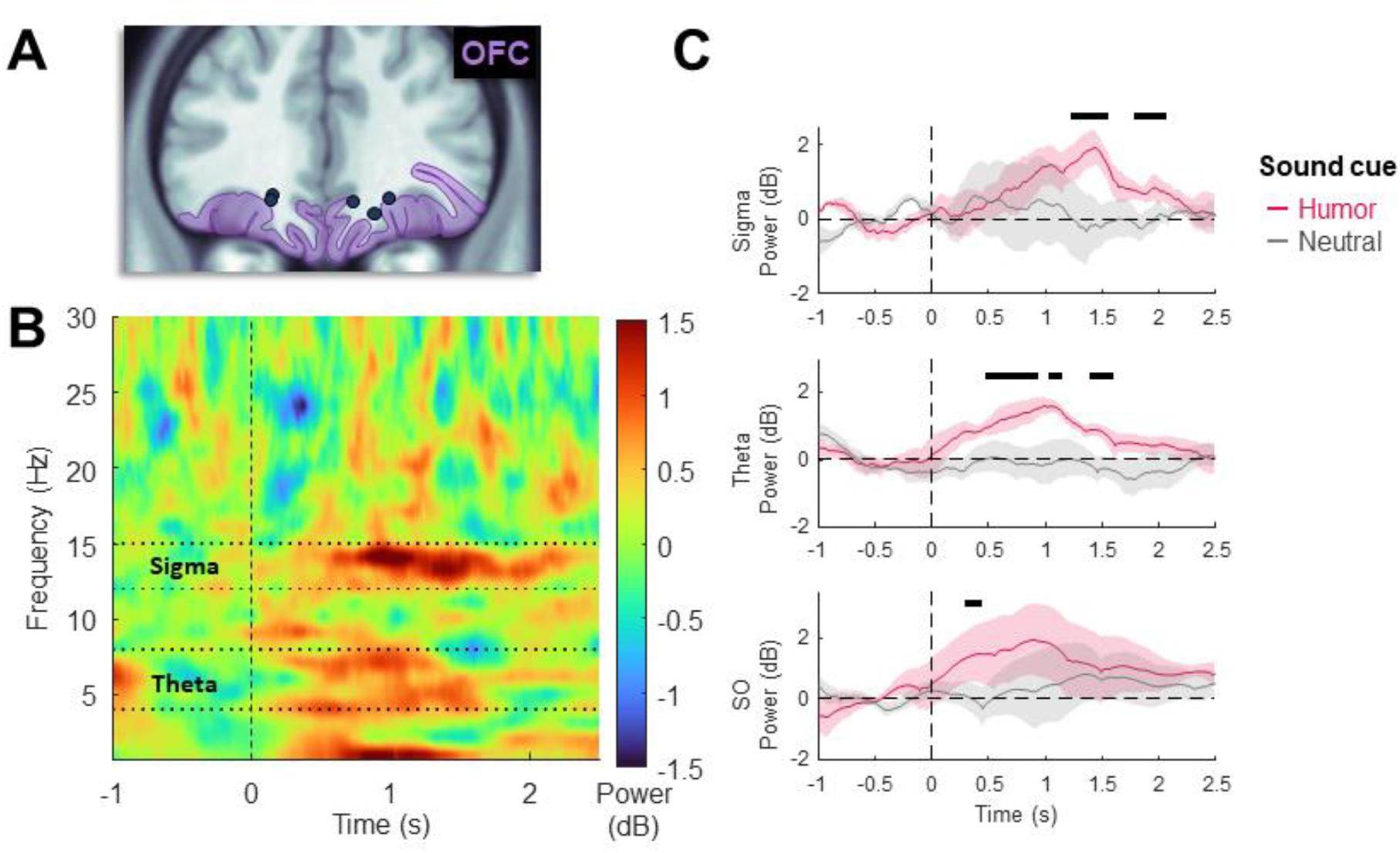
Neural responses to auditory cues during sleep. (A) Electrode position within the OFC (purple shading) on MNI152 template brain for each patient (dark blue dots). (B) Mean difference in time-frequency response to emotional versus neutral auditory cues in the OFC of the Sleep group. The heatmap depicts the amplitude (in color) of the response to stimulation at each frequency (y-axis) and time relative to stimulus onset (x-axis). (C) Typical frequency band of sleep research where investigated (i.e. SO 0.75Hz, theta 4-8Hz and sigma 12-15Hz). The detailed time courses of power change for these relevant frequency bands are plotted. The plain line corresponds to the average time course in response to neutral (gray) and emotional (fuchsia) cues across patients and the shaded area around the line to the standard error of the mean. Each frequency band was analyzed at different timings represented by light gray areas. Dark thick horizontal lines represent statistical differences between responses to emotional and neutral cues.

Next, we tested whether the sustained increase in theta activity during sleep oscillations (i.e. slow-oscillations and spindles) in OFC was linked with enhanced communication with other brain regions involved in emotion (amygdala) or memory (hippocampus). Specifically, we investigated whether emotionally-conditioned sounds (as compared to sounds associated with neutral pictures) boosted theta phase synchronization between these brain regions by computing the weighted phase-locking index (Vinck et al., 2011). We found greater connectivity of OFC with the hippocampus in the theta range (Fig. 3B right panel; largest cluster from 547 to 1130 ms; 0.042 ±0.028 dB; bootstrap resampling with N=5; z=3.815; p<0.001; CI=[0.021,0.065]; d=1.53), lasting over the entire largest window of increased theta oscillation for emotionally-conditioned sounds (see Fig. 2C for the extent in time of theta increase in the OFC). The same analysis did not show any significant difference for the wake group (Fig. 3B left panel), for amygdala-OFC connectivity or hippocampus-amygdala connectivity (Fig. S4), or for the sigma frequency range.

**Figure 3.**
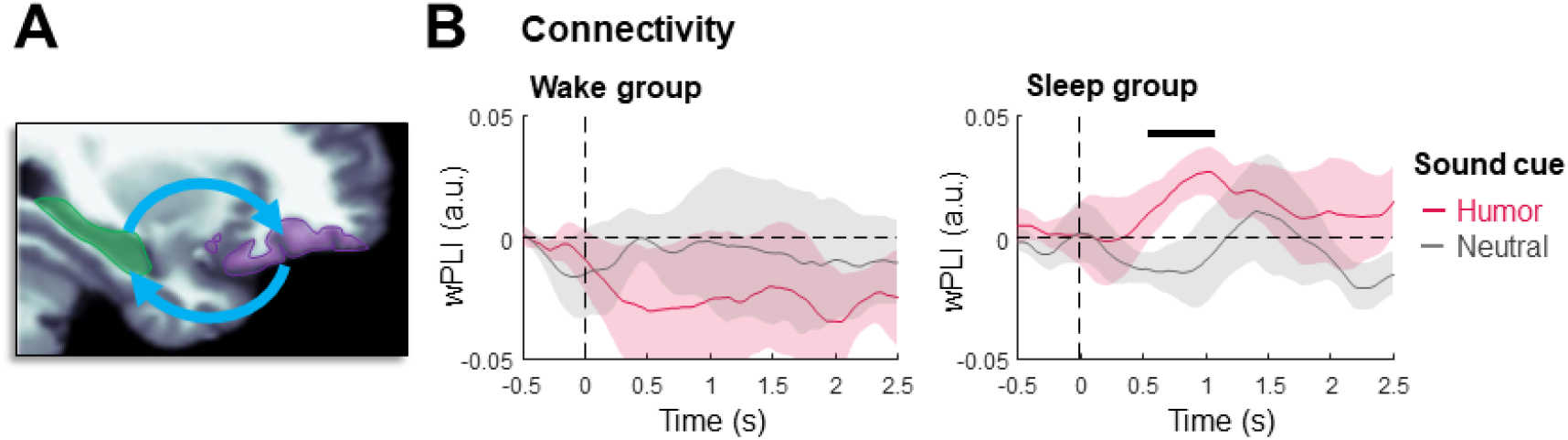
Phase synchrony between hippocampus and OFC cortex. (A) The hippocampus (green) and the OFC cortex (purple) are depicted on a slice of the MNI152 brain template. (B) Time course of the phase locking value after the presentation of the reward-conditioned (humor; fuchsia stroke) and the non-reward-conditioned (neutral; gray stroke) auditory cues. Dark thick horizontal lines represent statistically significant differences between the two conditions.

Finally, we explored the possibility that TMR applied during a daytime nap (compared to equivalent rest period of wakefulness) would differentially influence subsequent brain responses to the perception of emotional and neutral pictures. Overall, picture presentation induced an initial increase of all frequencies of the power spectrum in all three regions followed by a broad decrease of all frequencies of the power spectrum in the OFC (Fig. S5B top panel) and of frequencies below 15 Hz in the hippocampus (Fig. S5A top panel) and the amygdala (Fig. 4A and 4B). In the amygdala, the theta frequency range was rather spared by this broad decrease (Fig. 4C). We investigated the time courses of theta power, as this well-described rhythm is involved in memory encoding and retrieval (see above and Introduction section, see also Herweg et al., 2020), and compared them between the Sleep and Wake groups, in the three regions of interest, during picture presentation both at encoding and during the memory test. We did not find any close to significant differential group effect in the OFC (Fig. S5, middle and bottom panels). In the hippocampus, and in the amygdala, we found no initial group difference during encoding (Fig. 4D top panels, 4E and S5A middle panels; no significant difference at any timepoint). By contrast, during the memory test, theta power decreased more when seeing emotional than neutral pictures in the Sleep group both in the amygdala (Fig. 4D bottom panels; Cluster from 500 to 1970 ms; - 1.054±0.429 dB; Bootstrap resampling with N=6; z=-6.592; p<0.001; CI=[-1.384,-0.769]; d=-2.46) and in the hippocampus (Fig. S5 bottom panels; Cluster from 836 to 1580 ms; - 0.713±0.217 dB; Bootstrap resampling with N=6; z=-8.835; p<0.001; CI=[-0.878,-0.561]; d=-3.29), but not in the Wake group. In addition, in the amygdala of the Sleep group, the interaction between the experimental phase (encoding vs memory test) and the emotionality of pictures was significant (Cluster from 1160 to 1590 ms; 1.156±1.192 dB; N=6; z=2.603; p<0.001; CI=[0.444,2.116]; d=0.97).

**Figure 4.**
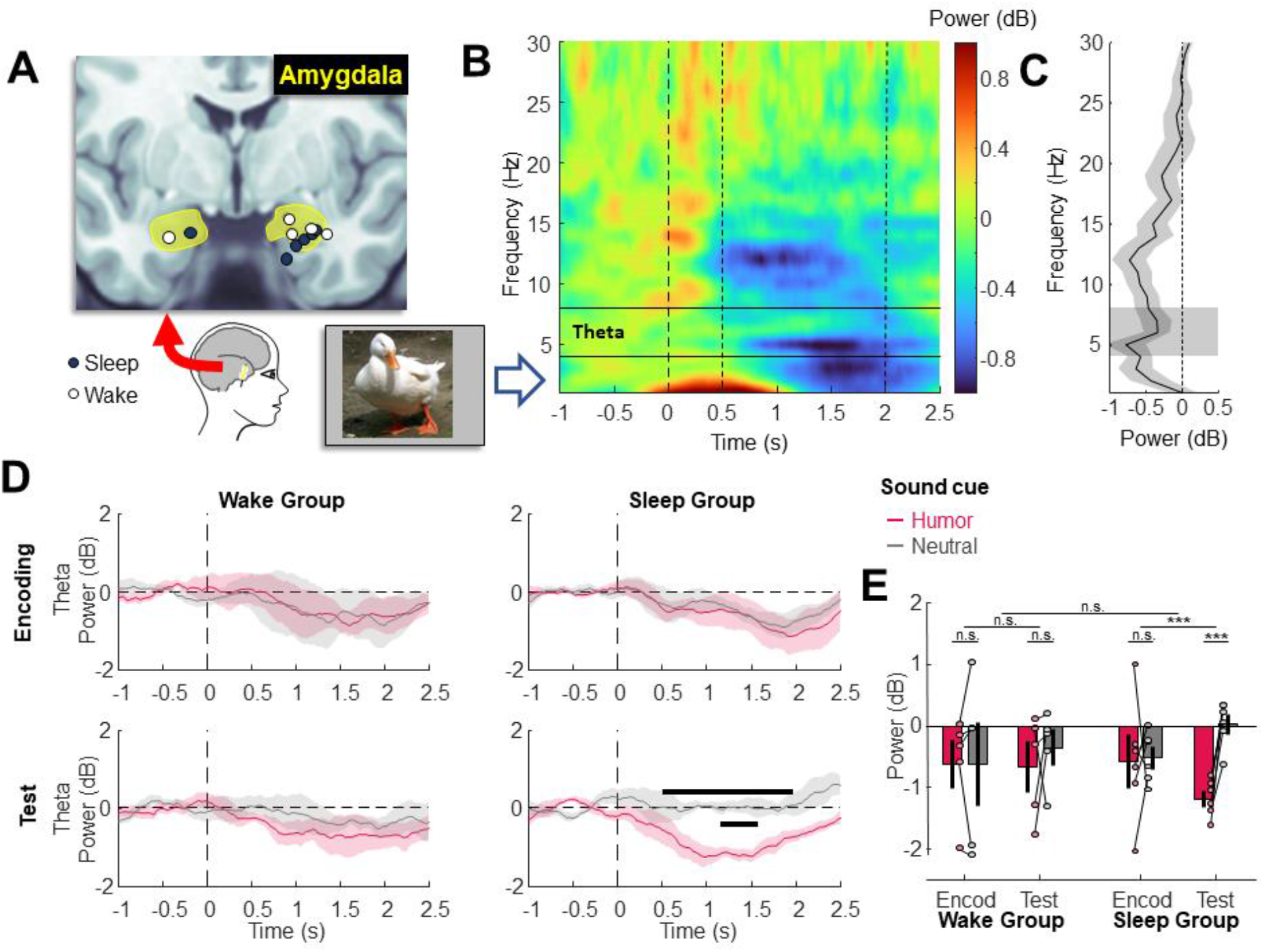
Changes in theta power during memory retrieval after nap with TMR. (A) Electrode positions within the amygdala (yellow shading) on MNI 152 template brain for each patient in the sleep (dark blue dots) and wake (white dots) group. (B) Averaged time-frequency response to pictures in both the encoding and memory test phases. The heatmap depicts the power changes of the amygdala signal relative to the baseline while patient saw pictures (neutral and funny together, encoding and test phase together ; see methods). Visual observation of the heatmap suggests that picture induce a broad increase of low (below 3Hz) and high (above 10Hz) frequencies power and then a decrease of power below 15Hz. (C) A visual observation of the average spectrum of the responses between 0.5 and 2 seconds post-stimulus onset reveals a rather spared theta frequency power within the global decrease of frequencies below 15 Hz. (D) The time course of theta power relative to picture onset was computed for the wake (left subpanels) and sleep (right subpanels) groups at encoding (top subpanels) and test (bottom subpanels) phase separately. Humorous pictures (fuchsia trace) induced a similar time course of theta power than neutral pictures (gray trace) in all conditions except at the test phase for the sleep group (bottom right subpanel), suggesting that the TMR phase during nap had an influence on memory traces of emotional pictures. Black thick horizontal line above zero represents significant difference of response to emotional and neutral pictures and black thick horizontal line below zero represents a significant interaction between emotionality of pictures (emotional versus neutral pictures) and experimental phase (encoding versus memory test). (E) The theta power was averaged for each individual and condition during the time points of significant interaction of emotionality x phase and displayed on the bar graph. Bars represent the mean across patients, small circles the individual data and the bold plain line at the center of the bar the standard error of the mean.

When focusing on the largest cluster of timepoints when the interaction between the emotionality of pictures and the experimental phase was significant, the difference between humorous and neutral pictures was significant for the Sleep group during the test phase (Fig. 4E; -1.212 ± 0.430 dB; bootstrap resampling with N=6; z=-7.560; p<0.001; CI=[-1.547,-0.925]; d=-2.82) but not for the Wake group (−0.314 ± 1.056 dB; bootstrap resampling with N=6; z=-0.743; p=0.391; CI=[-1.070,0.498]; d=-0.30). The interaction between the experimental phase and the emotionality of the pictures was not significant for the wake group (Mean difference±SD: 0.322±1.426 dB; bootstrap resampling with N=6; z=0.564; p=0.597; CI=[-0.657,1.605]; d=0.23) but significant for the sleep group (Mean difference±SD: 1.156±1.192 dB; bootstrap resampling with N=6; z=2.603; p<0.001; CI=[0.444,2.116]; d=0.97). There was no triple interaction between patient groups, experimental phase and emotionality of pictures however (Mean difference±SD: - 0.835±1.301 dB; bootstrap resampling with N=11; z=-0.014; p=0.389; CI=[0.793,-2.384]; d=-0.64).

## Discussion

Events associated with an emotion are better encoded in memory, as compared to emotionally-neutral events (LaBar & Cabeza, 2006). The emotional dimension (e.g. subjective experience, physiological expression, neuronal activity) of a recently encoded event might therefore constitute a key determinant for its subsequent consolidation. The reactivation of these emotional memory traces occurs spontaneously during sleep but can also be boosted by a Targeted Memory Reactivation (TMR) procedure. When a sensory cue reactivates the memory of an emotional situation, it may automatically reinstate its associated affective features (Cairney et al., 2014; Lehmann et al., 2016; Sterpenich et al., 2014). It is thought that, during sleep, TMR works through the reactivation of memories, but it is unclear whether (i) brain regions involved in emotion processing reacted more to these memory traces during sleep or if the affective dimension of the original event may also be reinstated across emotion/reward brain networks, and how (ii) affective and memory systems interact when emotional memories are reactivated in sleep.

By recording intracranial EEG signal from depth electrodes implanted in epileptic patients undergoing a presurgical investigation and using a TMR paradigm, we investigated the activity of key brain regions involved in emotion and memory processing during the reinstatement of emotional associations in sleep. Prior to sleep, we associated one sound with humorous pictures (i.e., emotional conditioning) and another sound with neutral pictures. When the patients were in stable NREM sleep, we found an initial increase in sleep oscillation activity, followed by a sustained increase in theta power, and a later increase in sigma power in OFC in response to the humor-conditioned sound cue (compared to the neutral sound cue, Fig. 2). The observed temporal organization between slow oscillations and sleep spindles (i.e., peak in sigma range) is reminiscent of the well-known slow-oscillation-spindle complex, whereby spindle generation increases during the up-state of slow oscillations (Clemens et al., 2007; Klinzing et al., 2016; Marshall et al., 2020; Staresina et al., 2015; Takeuchi et al., 2016). Our results are also consistent with those from a previous scalp EEG TMR study, in which Lehman et al. (2016) used a similar paradigm and reported that cues related to emotional memories selectively enhanced slow-waves activity and then sigma power. Moreover, we found a sustained theta activity, starting at about 500 ms post-stimulus onset (Fig. 2C), which also converges with the results from Lehmann et al. (2016). A similar theta power increase, at a slightly earlier latency, was observed in response to “hard-wired” emotional cues (i.e., angry voices; Blume et al., 2017). In the present study, we could establish that the theta increase caused by the presentation of emotionally-conditioned sounds during sleep likely originated partially from the OFC. Such an emotion-related modulation of theta power provides additional evidence supporting that the sleeping brain can detect whether a sound has acquired an emotional significance or not (Beh & Barratt, 1965; Blume et al., 2017, 2018; Lehmann et al., 2016; Perrin et al., 1999). Almost concomitant with the sustained theta boost, we observed an increase in the connectivity between the hippocampus and OFC also in the theta range (Fig. 3). One plausible interpretation is that the association between memory elements and their emotional value was reactivated by the auditory cue and expressed by an increased theta connectivity between memory (hippocampus) and emotion (OFC) networks. Such theta oscillations are believed to mediate a dialog between the hippocampus and cortical regions supporting memory retrieval during wakefulness (Herweg et al., 2016, 2020; Kaplan et al., 2014). Moreover, the sleep-related consolidation of emotional associations has been previously shown to strengthen the functional connectivity between the hippocampus and the prefrontal cortex (Sterpenich et al., 2007). Thus, our data offer an explanatory mechanism linking these separate observations by demonstrating that the reinstatement of emotional memories during NREM sleep involves a transient enhancement of theta power in the OFC together with a sustained increase in theta connectivity with the hippocampus.

After this communication between the hippocampus and the OFC, we also observed the emergence of spindles in the OFC. Several studies indicated that the prioritization of emotional memories (over neutral memories) for consolidation during sleep is associated with an increase of sleep spindles. Indeed, as spindles often follow a slow wave and are associated to brain plasticity, we can hypothesis that emotion boost the classical mechanism of brain consolidation during sleep (Cairney et al., 2014; P. Hu et al., 2006; Igloi et al., 2015; Sterpenich et al., 2007; Wagner et al., 2006). In addition, patients stimulated during sleep (i.e., Sleep group) showed a subsequent decrease of theta power selectively for cued emotional pictures during recall. Theta rhythm is often linked to memory encoding and retrieval. For instance, several scalp EEG and MEG studies reported that increased theta power during encoding indexed better subsequent memory recall (Herweg et al., 2020; Staudigl & Hanslmayr, 2013). By contrast, studies using intracranial EEG usually documented a decrease in theta power in the medial temporal lobe (Lega et al., 2012) and in other cortical populations (Fellner et al., 2019; Long et al., 2014) associated with memory success (see Herweg et al., 2020 for a review). Based on these previous observations, we suggest that memory reactivation during sleep leads to a reduction in theta power at wakefulness during subsequent retrieval that we observed in the present study. This points toward a strengthening and a more efficient memory traces for emotional items after TMR applied during sleep (please note however that we did not find any group effect on memory performance).

To summarize, our data suggests that emotional cues presented during sleep potentiated oscillatory activity (SO, theta and sigma activity) in the OFC, paralleled by an increase in functional connectivity between the hippocampus and OFC in the theta range. Respectively, these induced oscillatory events are consistent with the reinstatement of the emotional dimension of recently encoded memories across emotion brain networks, and with the emergence of a functional dialog between affective and memory systems when emotional memories are reactivated in sleep. Such oscillatory activities might provide a mechanism whereby memory traces for emotional items are prioritized for consolidation during sleep. This proposal is further supported by a subsequent theta decrease observed selectively for emotional (humorous) pictures in the Sleep group during memory recall. Note that our small sample size did not allow to formally test for sleep-related change in behavioral memory performance.

Some limitations might apply to our study. Firstly, we only used positive (humorous) stimuli while most studies in healthy participants used negative (e.g. fearful, aversive) stimuli, as negative emotions usually lead to stronger memory effects. Because the patients were undergoing a stressful medical procedure (i.e. presurgical intracranial recordings) and may often show signs of elevated anxiety, we chose not to present aversive stimuli, which may cause lasting psychological distress in those vulnerable patients, hamper emotion-specific TMR responses or, in the worst case, require the discontinuation of the experiment. Secondly, regarding the OFC signal, we managed to record neuronal population from the ventral side of the prefrontal cortex (i.e. OFC and ventro-medial prefrontal cortex) but surrounding neural structures, in particular the nucleus accumbens and lateral prefrontal cortex, might have also contributed some activity in a few patients. Thirdly, more studies are needed to clarify the exact mechanisms underlying the changes of theta power during memory recall. In another study, two types of theta rhythms, with distinct frequencies, were observed in the hippocampus and other cortical populations and the authors postulated that they represent different processes of memory encoding and retrieval (Lega et al., 2012). In addition, a ‘spectral tilt’, which supposedly reflects a change in the neural excitatory/inhibitory balance, was suggested to possibly mask increases of theta range activity (Donoghue et al., 2020; Herweg et al., 2020). Our data suggests a broad decrease of low-frequencies after picture viewing consistent with a ‘spectral tilt’ but the decrease of theta power following humoristic picture viewing after the nap could either reflect a larger tilt of the spectrum or a specific decrease in theta rhythm related to the strength of memory traces (Fellner et al., 2019). Finally, it is unsure whether our manipulation of consolidation processes with TMR induced the observed change of evoked theta power during memory recall or whether mere sleep exposure was sufficient to produce these effects. The literature on mental fatigue shows that stimuli-evoked theta power is influenced by mental fatigue accumulated over the day (Wascher et al., 2014; Yu et al., 2021). As epilepsy monitoring is an exhausting procedure and our participants were probably tired and sleep deprived, the nap of the sleep group might, by itself, have restored the ability to modulate brain oscillations in response to stimuli without the intervention of TMR.

We were also not able to demonstrate the causality of the brain oscillations during sleep on memory performances on the next due to the large variability of memory abilities in this type of patients. Finally, we did not record brain oscillations in the ventral tegmentum area or in the striatum, two key regions involved in the reward processing and that has been shown to dialogue with the hippocampus during sleep (Pennartz et al., 2004; Lansink et al., 2009). No patient had electrodes in these regions which play a major role in the above-mentioned processes.

Our results show that, beyond the reinstatement of hippocampal and related cortical sensory representations of a memory episode (Klinzing et al., 2019), cued reactivation in sleep encompasses networks mediating other critical dimensions of the memory episode, such as its emotional relevance. Emotional reinstatement was also accompanied by changes in oscillatory activities typical of memory consolidation mechanisms, including an increase functional dialog between the hippocampus and OFC (i.e., increased theta phase-synchrony).These findings also support that the reprocessing of rewarded memories during sleep implicates sleep-related oscillatory mechanisms enabling a functional dialog between reward brain regions and the hippocampus. We suggest that these changes in oscillatory activity might be implicated in the prioritization of emotional memory consolidation during sleep.

