## Supplementary material for "Reinstatement of emotional associations during human sleep: an intracranial EEG study"

| | Wake Group | | Sleep Group | | All | |
|---|---|---|---|---|---|---|
| Measure | Emo | Neu | Emo | Neu | Emo | Neu |
| Rating - Encoding | 1.55±0.62 | 0.32±0.36 | 1.62±0.47 | 0.31±0.40 | 1.59±0.52 | 0.32±0.37 |
| Hits (%) - Memory test | 92.33±5.96 | 82.00±9.82 | 82.07±21.37 | 65.91±31.13 | 86.74±16.47 | 73.22±24.36 |
| Recognized (%) - Memory test | 77.33±17.42 | 62.33±22.69 | 70.59±27.55 | 48.08±32.63 | 73.66±22.65 | 54.56±28.17 |
| Familiar (%) - Memory test | 15.00±12.58 | 19.67±14.88 | 11.48±8.79 | 17.83±14.46 | 13.08±10.26 | 18.66±13.93 |
| Correct rejections (%) - Memory test | 76.00±25.76 | 69.33±25.97 | 86.04±9.22 | 75.56±16.95 | 81.47±18.31 | 72.73±20.59 |

**Table S1**. Behavioral performance during encoding and memory test. Average humor rating of emotional and neutral pictures (mean±SD) during encoding is presented in the first row. During memory test, percent of hits (i.e., recognized and familiar together), recognized, and familiar pictures among pictures already presented during encoding are shown in the second, third and fourth rows respectively. Percent of correct rejections of pictures never seen during encoding are shown in the fifth row.

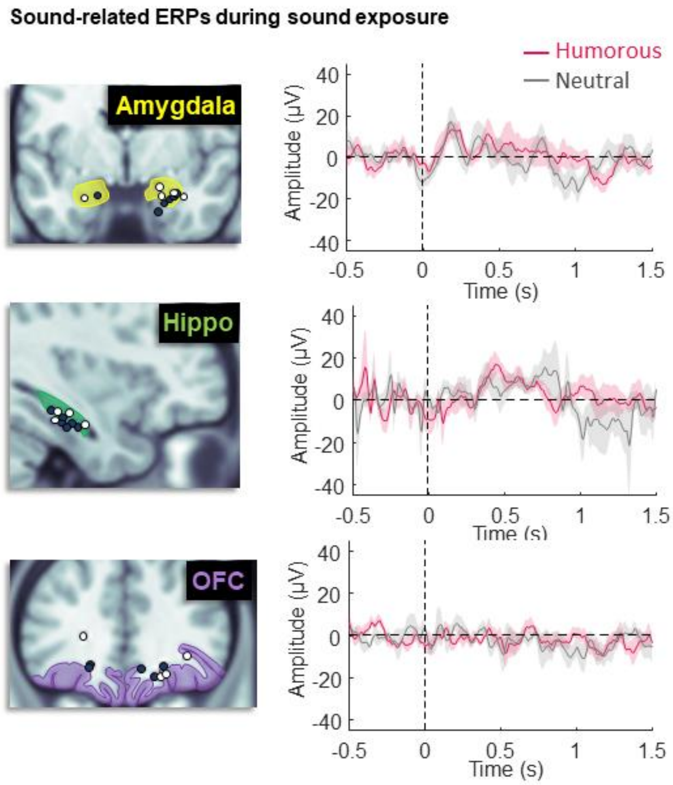


**Figure S1**. ERPs to sounds during the sound exposure. For each brain region depicted on the left, the ERPs across participants was computed for the to be emotional (fuchsia) and to be neutral (gray) auditory cues. Note that the small number of trials per condition in this part of the experiment might explain the noisy signals.

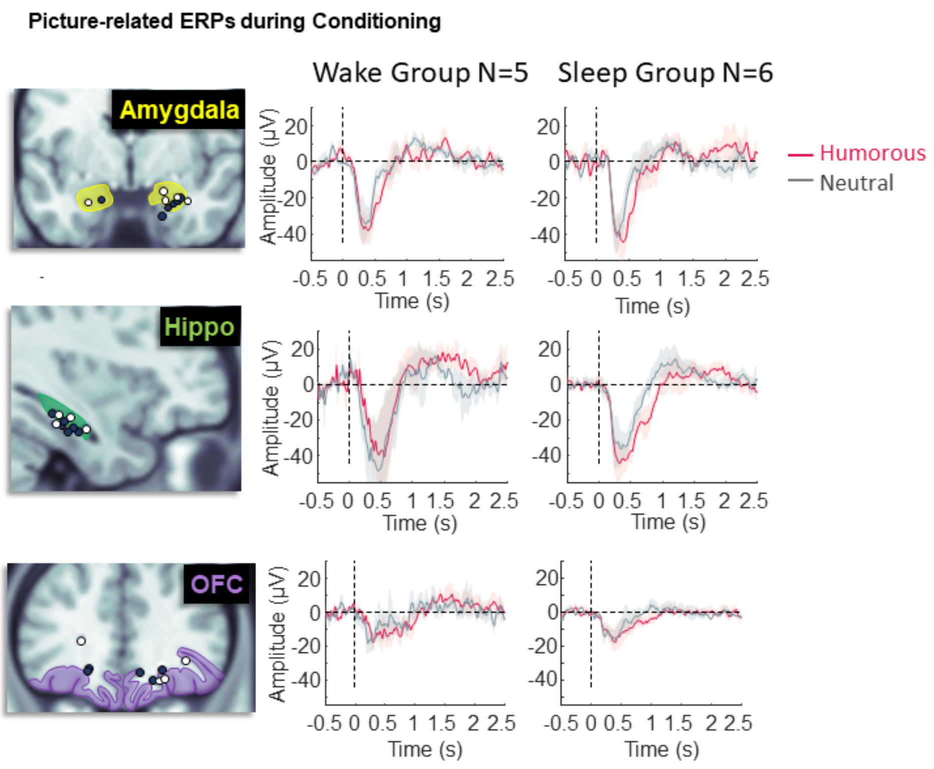


**Figure S2**. ERPs to pictures during conditioning. For each brain region depicted on the left, the ERPs across participants was computed for emotional (fuchsia) and neutral (gray) pictures.

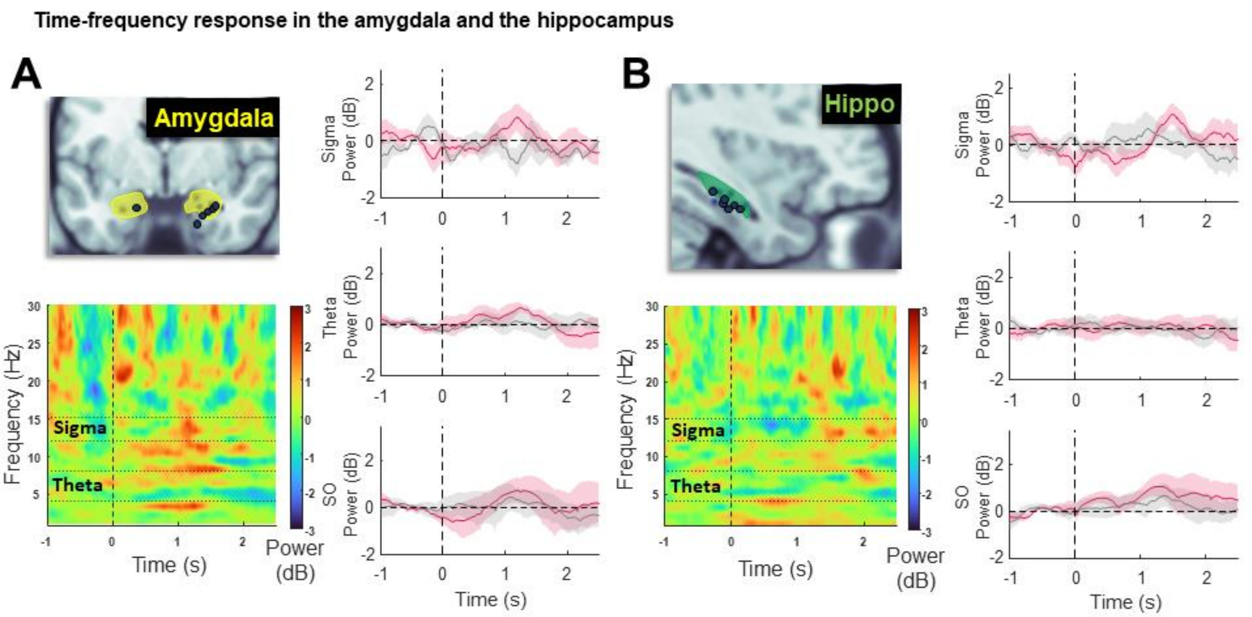


**Figure S3**. Time-frequency responses to auditory cues of the sleep group during the TMR session in (A) the amygdala and (B) the hippocampus. Each panel depicts the electrode positions in each region (top left subpanel), the heatmap of the difference of time-frequency response (bottom left subpanel) and the responses in canonical frequency ranges (right subpanels; from top to bottom: sigma, theta and SO power respectively) for emotional (fuchsia) and neutral (gray) auditory cues.

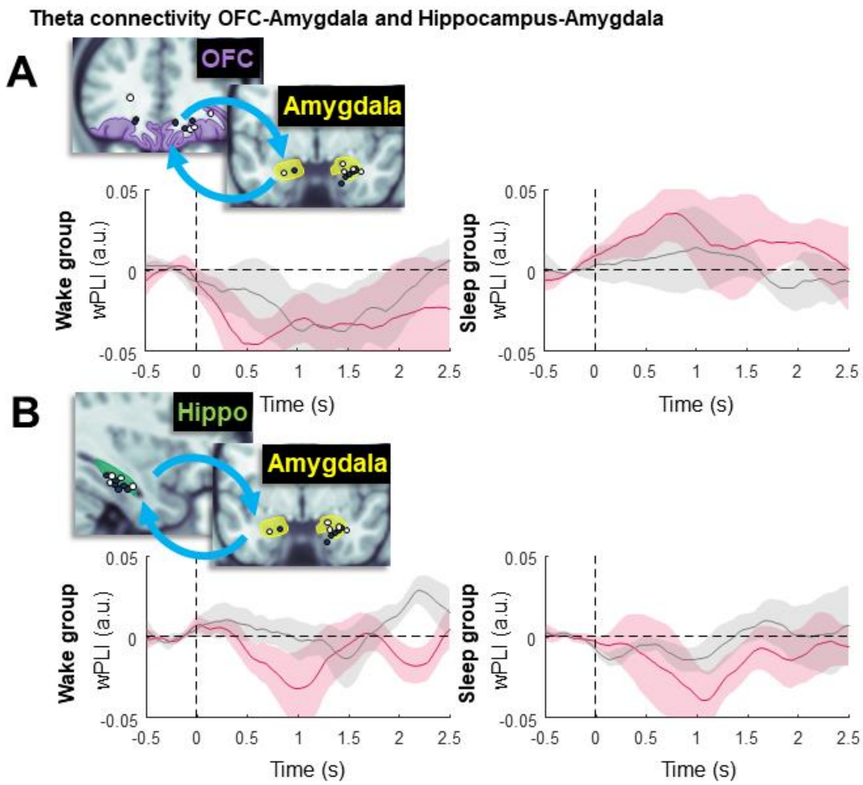


**Figure S4**. Theta connectivity in response to auditory cues during the TMR session between the OFC and the amygdala (A) and between the hippocampus and the amygdala (B). Each panel depicts the regions implicated in the connectivity (top left corner) and the time courses of weighted phase-locking index in response to emotional (fuchsia) and neutral (gray) cues in the wake group (left subpanel) and sleep group (right subpanel).

Time-frequency response to pictures with a focus on theta power in the hippocampus and the OFC

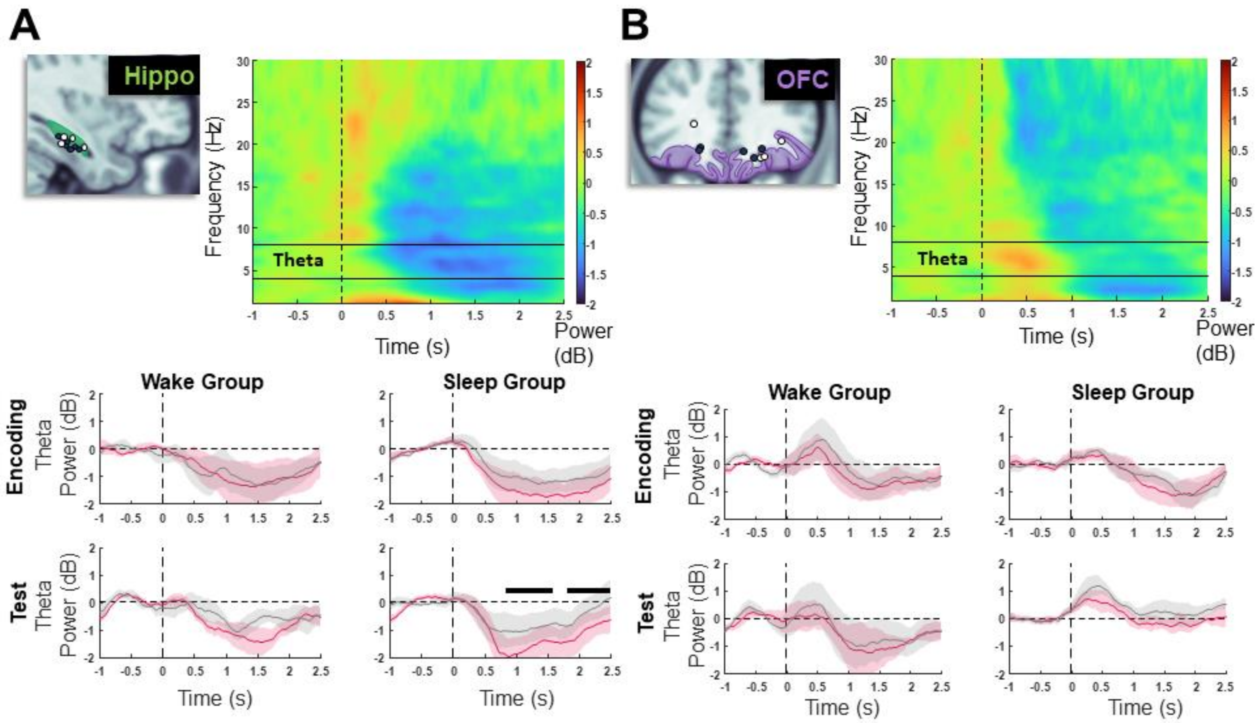


**Figure S5**. Modulation of theta power in response to emotional and neutral pictures between encoding and memory test. Similarly to Figure 4, time-frequency responses to all pictures (top right subpanel; emotional and neutral pictures together; during encoding and memory test) were computed in the hippocampus (A) and OFC (B). The time course of theta power was computed for the wake and sleep group separately (left and right subpanel respectively), during encoding and memory test separately (middle and bottom subpanel respectively) for emotional pictures (fuchsia) and neutral pictures (gray). Black thick horizontal lines represent a significant difference of response to emotional and neutral pictures. Note that only the hippocampus (A) shows a difference of response in the theta frequency range to emotional pictures during memory test. No significant interaction between emotionality of pictures (emotional versus neutral) and phase of the experiment (encoding versus memory test) was found.
